# Fecal transplant allows transmission of the gut microbiota in honey bees

**DOI:** 10.1101/2023.11.29.569223

**Authors:** Amélie Cabirol, Audam Chhun, Joanito Liberti, Lucie Kesner, Nicolas Neuschwander, Yolanda Schaerli, Philipp Engel

## Abstract

The gut of honey bees is colonized by symbiotic bacteria during the first days of adult life, once bees have emerged from their wax cells. Within five days, the gut microbiota becomes remarkably stable and consistent across individual bees. Yet, the modes of acquisition and transmission of the gut microbiota are to be confirmed. Few studies suggested bees could be colonized via contact with fecal matter in the hive and via social interactions. However, the composition of the fecal microbiota is still unknown. It is particularly unclear whether all bacterial species can be found viable in the feces and can therefore be transmitted to newborn nestmates. Using 16s rRNA gene amplicon sequencing we revealed that the composition of the honey bee fecal microbiota is strikingly similar to the microbiota of entire guts. We found that fecal transplantation resulted in gut microbial communities largely similar to those obtained from feeding gut homogenates. Our study shows that fecal sampling and transplantation are viable tools for the longitudinal analysis of bacterial community composition and host-microbe interactions. Our results also imply that contact of young bees with fecal matter in the hive is a plausible route for the acquisition of the core gut microbiota.

## Introduction

Over the past decade, honey bees (*Apis mellifera*) have become pivotal insect models for the study of gut microbiota evolution and function^1–3^. This is due to the relatively simple composition and consistency of their gut microbiota, the possibility to study *in-vitro* and *in-vivo* defined communities of gut bacteria, as well as the recent opportunity to genetically engineer some of the gut symbionts^4–6^. The honey bee gut microbiota has also attracted a lot of attention due to its important role in shaping the health and behavior of these essential pollinators^7–10^.

The honey bee gut is subdivided into four distinct sections: the crop and midgut contain few bacteria, while the ileum and rectum, together forming the hindgut, contain most core members of the honey bee gut microbiota in different proportions^4,11^. The core bacteria *Gilliamella* and *Snodgrassella* are predominant in the ileum, where they form a biofilm, while *Bombilactobacillus* Firm-4, *Lactobacillus* Firm-5, and *Bifidobacterium* dominate the rectum community^11–13^. How such stable gut bacterial communities are transmitted between individuals remains unclear in this social insect.

Honey bee workers are known to progressively acquire their gut microbiota during the first week of adult life in the hive, after emerging from their wax cells^11,12^. The presence of adult nurse bees^11^ or fresh pollen from the hive^14^ in the environment of newly emerged bees was shown to promote the acquisition of the core microbiota. Suggested mechanisms in these studies are: (*i*) direct transmission *via* trophallaxis behavior, where bees actively exchange the food content of their crop in a mouth-to-mouth interaction, and (*ii*) indirect transmission *via* contact with the fecal matter of nurse bees deposited in the hive environment. Recent studies found that trophallaxis with nurse bees alone was not sufficient^12^ and even unnecessary^14^ for newly emerged bees to acquire the core gut microbiota. Instead, exposure to hindgut homogenate successfully led to a gut microbiota community similar to the one of hive bees. Gut homogenates, however, not only contain fecal matter, but also the communities of bacteria attached to the gut epithelium. The source of gut microbiota transmission thus remains ambiguous. Since honey bees do not systematically defecate in laboratory conditions while kept in cages, the use of hindgut homogenates over isolated fecal matter has so far been predominant in the field; whether it is to investigate the mechanisms underlying microbiota transmission or to inoculate microbiota-free (MF) individuals in the context of *in vivo* experiments. Nonetheless, work carried out by our group and others established protocols for routine feces sampling of honey and bumble bees, respectively^5,15,16^. It remains uncertain whether all gut microbiota phylotypes, especially those preferentially colonizing the ileum and forming biofilms, are viable and present in sufficient quantities in fecal matter to allow microbiota transmission across individuals.

Thus, our investigation set out to validate the hypothesis that the honey bee gut microbiota can be naturally transmitted through contact with fecal matter by quantifying the relative transmission of the different bacteria present in feces. Using qPCR quantification and amplicon sequencing targeting the 16S rRNA gene, we compared the bacterial taxonomic composition in the feces and guts collected from the same nurse bees (generation no. 1) to understand whether the feces of honey bees provide a robust proxy for their gut microbiota (**Fig. 1**). We then analyzed the bacterial taxonomic composition in the gut of bees fed with feces or gut homogenate a week post-inoculation to determine whether ingestion of feces allows transmission of the microbiota from adults to newly emerged microbiota-free bees (generation no. 2). Our results demonstrate that the gut microbiota composition can be non-invasively monitored using fecal sampling, and that transplantation of fecal matter into microbiota-free bees is a reliable and ecologically relevant method to study microbiota transmission and host-microbe interaction.

**Figure 1.**
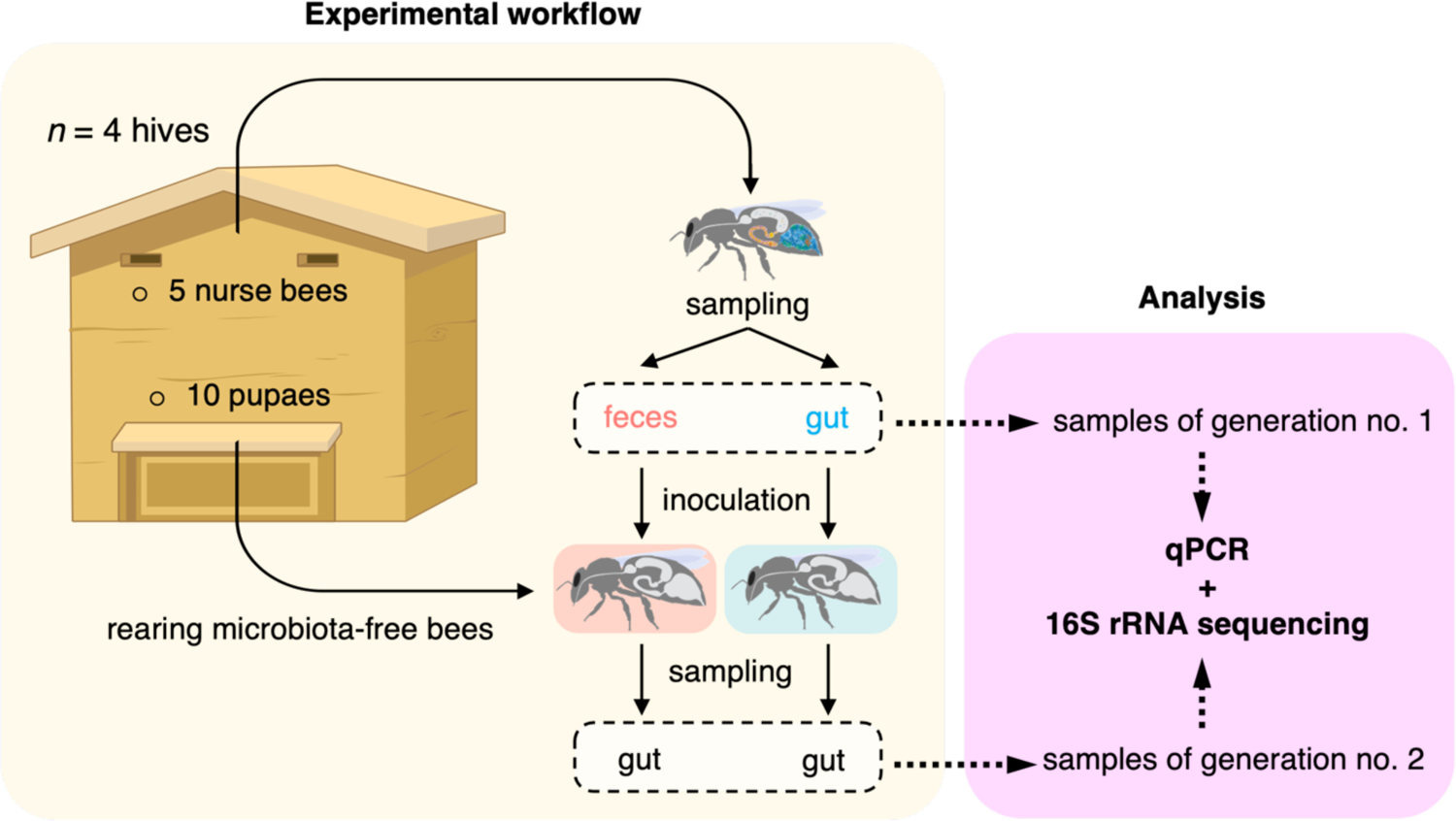
Schematic outline of the experimental workflow. The feces and gut from five nurse bees were collected to compare their bacterial composition (generation no. 1) and to inoculate five microbiota-free newly emerged bees (generation no. 2). A week post-inoculation, the guts of inoculated bees were collected, and their bacterial composition assessed. Bacterial total and relative abundances in the feces and gut samples were measured by quantitative PCR and 16s rRNA gene amplicon sequencing, respectively. The experiment was replicated four times using distinct hives.

## Results

### Characterization of the honey bee fecal microbiota

To establish whether feces of honey bees are a robust proxy for their gut microbiota, we compared the microbial communities present in feces *versus* gut samples of nurse honey bees from four distinct hives (Fig. 2). Honey bee feces were rich in bacteria, with a median bacterial load of 1.58·10^6^ cells μl^-1^ of feces (95% CI [9.20·10^5^, 2.59·10^6^]) (**Supplementary Fig. 1**). More importantly, the bacterial communities present in feces were remarkably similar to the ones found in the guts of naturally colonized honey bees (Fig. 2). The predominant genera of the gut microbiota of honey bees were detected in both gut and fecal samples, namely *Bombilactobacillus* Firm-4, *Lactobacillus* Firm-5, *Gilliamella*, *Snodgrassella*, *Bifidobacterium*, *Frischella, Bartonella, Commensalibacter* and *Apilactobacillus* (formerly *Lactobacillus kunkeei*) (Fig. 2a) ^1,17,18^. This was the case for all samples across the different hives tested, with the exception however of two bees from hive 15, which appeared to have very low bacterial complexity. We considered these samples as outliers that may have arisen from technical errors considering further analysis discussed below.

**Figure 2.**
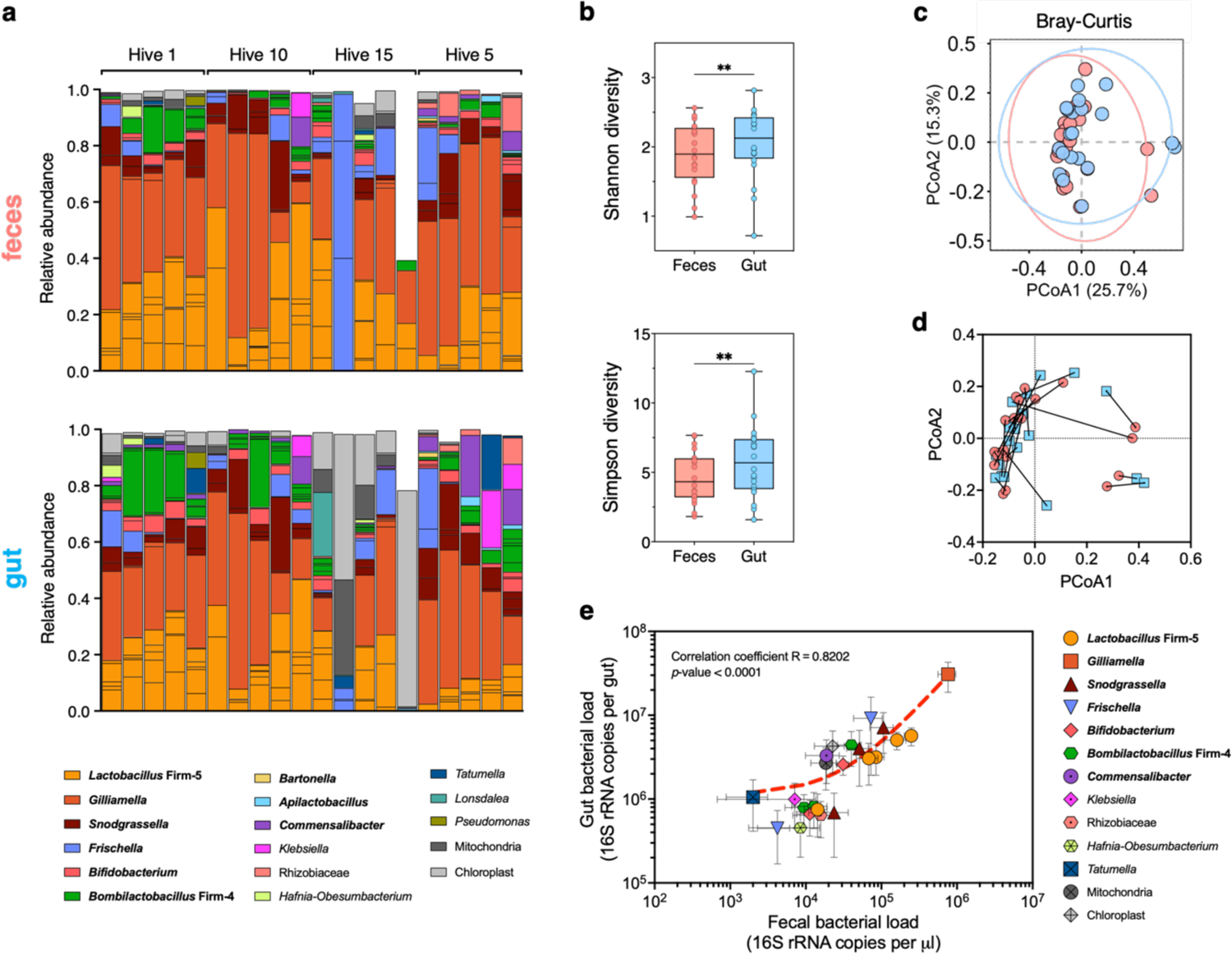
The fecal microbiota of honey bees is a robust proxy for their gut bacterial communities. **a** Stacked bar plots showing the relative abundance of identified amplicon-sequence variants (ASVs) grouped at the genus level in the feces (top panel) and gut tissues (bottom panel) of hive bees (generation no. 1). Vertically aligned bars represent samples sourced from the same individual. Their hive numbers are indicated. Only ASVs with relative abundance above 1% in at least 2 samples are displayed. Prevalent members of the honey bee gut microbiota are in bold. **b** Bacterial ⍺-diversity was significantly higher in the gut than in the feces of hive bees according to both the Shannon (Wilcoxon matched-pairs test (two-tailed)) and Simpson indexes (paired t-test). ** P < 0.005. **c** Principal-coordinate analysis (PCoA) based on the Bray-Curtis dissimilarity index showed no significant difference in β-diversity between the feces (red) and gut (blue) samples (PERMANOVA test, not significant). **d** Procrustes analysis of relative ASV abundances in the feces (red) against gut (blue) samples of hive bees revealed a significant agreement of comparison. Longer lines on Procrustes plots indicate more dissimilarity between samples sourced from the same individual. **e** Scatter plot showing a significant correlation between the absolute abundances of bacteria genera in the gut and the feces samples (Pearson’s correlation coefficient and *p-*value are displayed). Only ASVs with absolute abundance above 1% in at least 5 samples are displayed for clarity. The red dotted line represents the linear regression curve (appearing non-linear due to log axes).

Diversity of the gut and fecal bacterial communities appeared overall comparable, as measured by alpha- and beta-diversity metrics. Alpha-diversity, which considers species richness and evenness within samples, was significantly higher in the gut samples compared to the fecal samples as measured using the Shannon index (Wilcoxon matched-pairs test, Z = 179, *p*-value = 0.0042) and Simpson metric (paired *t*-test, *t*_(19)_ = 3.39, *p*-value = 0.0031;Fig. 2b). A differential analysis revealed that only chloroplasts differed significantly in relative abundance between the fecal and gut samples likely because gut samples contained more pollen material (**Supplementary Fig. 2**; 13,711,506-fold change, adjusted *p*-value < 0.0001). Yet, the significant difference in alpha-diversity metrics remained after removing chloroplast DNA from the analysis (Shannon index: Wilcoxon matched-pairs test, Z = 177, *p*-value = 0.0056; Simpson metric: paired t-test, t_(19)_ = 2.7551, *p*-value = 0.0126). This difference was expected as feces constitute a subset of the gut samples. However, there was no significant difference between the microbiota structure of gut and fecal samples (PERMANOVA test based on Bray-Curtis dissimilarities, *p*-value = 0.1; Fig. 2c). Interestingly, Bray-Curtis dissimilarity matrices of the fecal and gut samples were positively correlated (Mantel test, *r* = 0.5, *p*-value = 0.0041). Consistently, a Procrustes analysis revealed a significant concordance between the feces and gut datasets (Fig. 2d; Procrustes randomization test, m^2^ = 0.45, p-value = 0.0060) indicating that fecal samples were on average more similar to the gut samples collected from the same individuals than to gut samples belonging to different individuals.

Finally, we observed a strong positive correlation in the absolute abundance of the most prevalent taxonomic groups between the gut and fecal samples, confirming that the gut colonization level of a given amplicon-sequence variant (ASV) was reflected by its concentration in the feces (Pearson correlation coefficient R = 0.82, *p*-value < 0.0001; Fig. 2e). Taken together, our results demonstrate that feces provide a robust proxy for the honey bee gut microbiota composition. Fecal samples allow to infer both the community membership (*i.e.* presence/absence of a bacteria) as well as to estimate the absolute bacterial abundances (*i.e.* levels of gut colonization) in the gut of individual bees.

### Transmission of the gut microbiota to microbiota-free honey bees *via* fecal transplantation

We next tested if ingestion of feces would be sufficient and equivalent to gut homogenates for microbiota transmission to newly emerged bees (Fig. 3). Five μl of fecal inoculum (7.89·10^5^ cells in the inoculum; 95% CI [4.60·10^5^, 1.29·10^6^]) was sufficient to successfully seed the gut of MF honey bees, resulting in colonization levels similar to the ones obtained when feeding 5 μl of gut homogenate (3.50·10^4^ cells in the inoculum; 95% CI [1.61·10^4^, 4.25·10^4^]; Wilcoxon matched-pairs test, Z = 26.00, *p*-value = 0.6226; Fig. 3a). The microbial communities in feces- and gut-inoculated bees reached a median of 1.54·10^8^ (95% CI [8.57·10^7^, 1.63·10^8^]) and 1.81·10^8^ (95% CI [1.03·10^8^, 2.22·10^8^]) cells per gut at day 7 post colonization, respectively. Additionally, the relative abundances of bacterial genera in those communities were again remarkably similar, with all prevalent genera of the bee microbiota found in the gastrointestinal tracts of individuals fed with gut or fecal inoculums (Fig. 3b). Even the bees that received fecal and gut inoculums sourced from the individuals of generation no. 1 from hive 15 that appeared to have a remarkably low-diversity microbiota (Fig. 2a), harbored a normal gut bacterial community here (Fig. 3a). This suggests that some technical issues may have distorted the gut community profiles of those individuals of generation no. 1 in our previous analysis. Alpha-diversity in the gut, measured with the Shannon and Simpson indexes, did not differ significantly between inoculum types (Fig. 3c; paired *t*-test, Shannon: *t*_(18)_ = −1.93, *p*-value = 0.07; Simpson: *t*_(18)_ = −1.45, *p*-value = 0.16), although there was a trend towards higher diversity in feces-inoculated bees. There was also no difference in community structure between bees fed the two different inoculum types (PERMANOVA using Bray-Curtis dissimilarities calculated from a matrix of absolute ASV abundance, *p*-value = 0.057; Fig. 3d). Honey bees inoculated with fecal material had on average slightly increased relative abundances of Bifidobacteria and Lactobacilli, which are rectum associated bacteria, than bees inoculated with gut homogenates (**Supplementary Fig. 3**). Yet, we found a robust positive correlation in the absolute abundance of the most prevalent genera composing the gut microbiota between honey bees fed with either gut or fecal inoculums (Pearson correlation coefficient R = 0.89, *p*-value < 0.0001; Fig. 3e). This confirms that feces are a good inoculum source, leading to a gut microbiota composition highly comparable to the one of bees inoculated with a gut homogenate.

**Figure 3.**
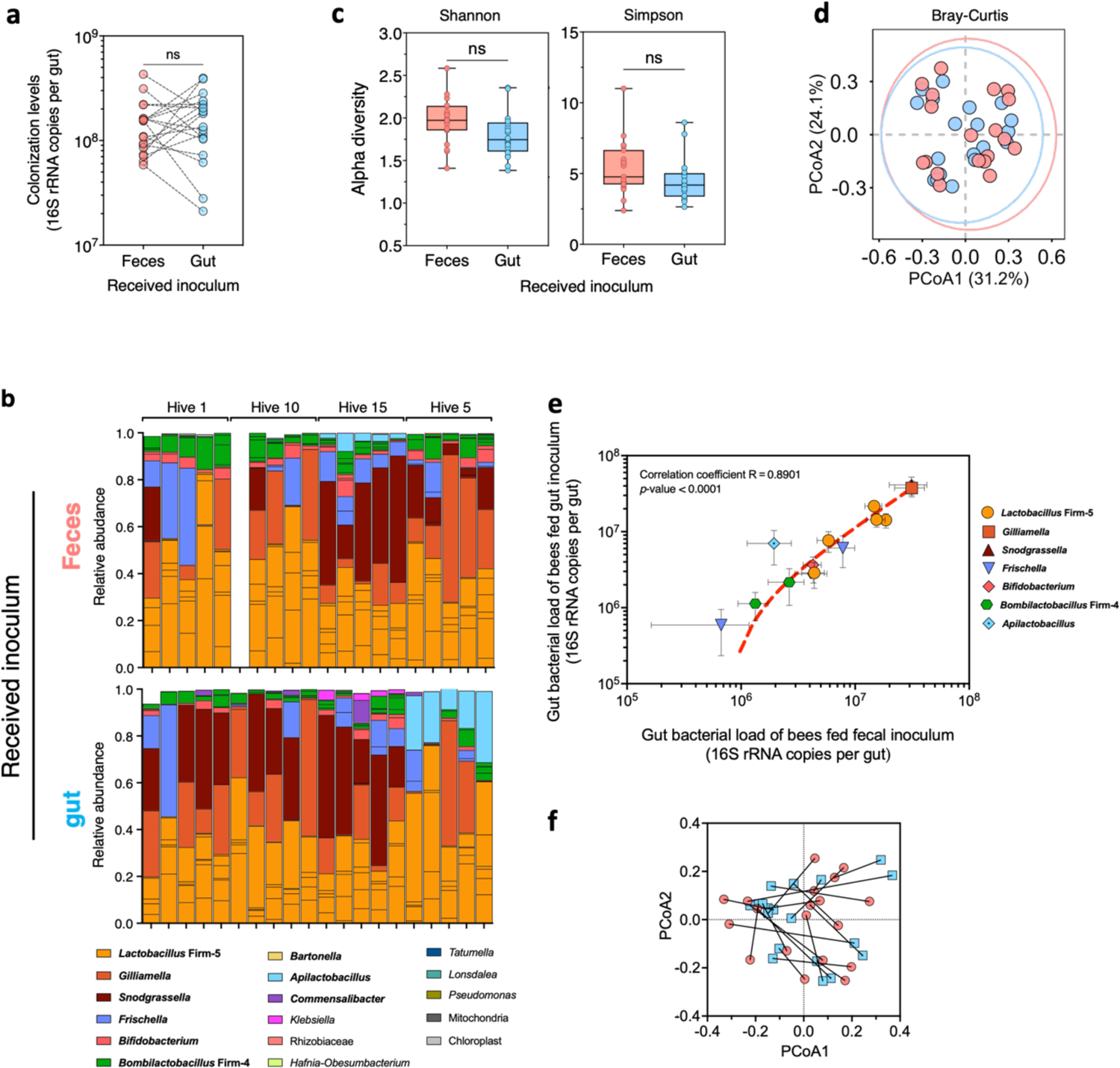
Fecal transplant allows transmission of the honey bee gut microbiota. **a** Colonization levels of bacteria in the guts of bees inoculated with either a gut homogenate or an aliquot of feces (generation no. 2) did not differ significantly (Wilcoxon matched-pairs rank test, not significant (ns)). Matching samples (*i.e.*, inoculums sourced from the same individuals) are connected by dotted lines. **b** Stacked bar plots showing the relative abundance of amplicon-sequence variants (ASVs), colored by their genus level classification, identified in the gut of bees inoculated with either feces (top panel) or gut homogenates (bottom panel). Vertically aligned bars represent matching samples. Their hive of origin is indicated above. For ease of visualization, only ASVs with a relative abundance above 1% in at least 2 samples are displayed. Prevalent members of the honey bee gut microbiota are shown in bold. One sample from hive ten was lost during the DNA extraction process. **c** Bacterial ⍺-diversity in the gut did not differ significantly between gut-inoculated and feces-inoculated bees based on both the Shannon and Simpson indexes (Unpaired *t*-tests (two-tailed), not significant, ns). **d** Principal-coordinate analysis (PCoA) based on the Bray-Curtis dissimilarity index showed no significant difference in β-diversity between the gut samples of feces-inoculated (red) and gut-inoculated (blue) bees (PERMANOVA test, not significant). **e** Scatter plot showing a significant correlation in absolute abundances of identified ASVs in the gut between feces-inoculated and gut-inoculated bees (Pearson’s correlation coefficient and *p-*value are shown within the plot). Only ASVs with an absolute abundance above 1% in at least 5 samples are displayed for clarity. The red dotted line represents the linear regression curve. **f** Procrustes analysis of relative ASVs abundances in the gut of feces-inoculated bees (red) against gut-inoculated bees (blue) revealed no significant agreement between sample pairs obtained from bees of generation no. 2 inoculated with feces and guts collected from the same donor. Longer lines on Procrustes plots indicate more dissimilarity between matching samples.

Finally, we performed two additional comparisons to assess the level of similarity in the microbiota that established in the gut bees of generation no. 2 and their respective donors in generation no. 1. First, we tested whether pairs of bees of generation no. 2 inoculated with feces and guts collected from the same donor were more similar between them than to other pairs of generation no. 2. Second, we tested whether the composition of the microbiota established in bees of generation no. 2 was more similar to that of the matched donor bees than to that of other bees of generation no. 1, for both feces and gut-inoculated bees independently. Pairs of bees of generation no. 2 inoculated with feces or gut homogenates originating from the same donor bees, were not more similar in gut microbiota composition than other generation no. 2 pairs (Fig. 3f; Procrustes randomization test, m^2^ = 0.66, *p*-value = 0.49; Mantel test, *r* = 0.0032, *p*-value = 0.47). The lack of similarity between matched pairs was further confirmed when comparing samples across generations (**Supplementary Fig. 4**). There was no significant concordance in the microbiota of donor and receiver bees across the two generations for both the feces (Procrustes randomization test, m^2^ = 0.62, *p*-value = 0.15; Mantel test, *r* = −0.17, *p*-value = 0.91) and the gut homogenate-inoculated bees (Procrustes randomization test, m^2^ = 0.65, *p*-value = 0.43; Mantel test, *r* = 0.06, *p*-value = 0.33). The absence of concordance between the community structures observed across generations for matched pairs suggests that community assembly is influenced by other factors distinct from the inoculum source.

## Discussion

Here, we characterized the bacterial composition of honey bee feces and found that the fecal microbiota resembles the gut microbiota. Moreover, inoculation of a small volume of feces to MF bees allowed all core microbiota members to establish in the gut, in similar relative and absolute amounts as the ones found in bees inoculated with a gut homogenate. The analysis of the fecal microbiota is commonly used in humans, laboratory rodents, and wild vertebrates to establish correlations between environmental factors, the gut microbiota, and host physiology^19,20^. However, the use of fecal matter as a proxy for gut microbiota composition in humans has been questioned, as the fecal microbiota was found to differ from the mucosa-associated microbiota^21–23^. By contrast, we found that honey bees collected from different hives at the nursing age harbored all core and most prevalent members of the gut microbiota in their feces, in proportions similar to those found in entire guts. Strikingly, bacteria known to colonize the fore part of the gut, namely the ileum, were also detected in the feces. This was unlikely due to the shedding and elimination of dead bacteria as these bacteria were viable and successfully established in the gut of feces-inoculated bees. As the honey bee gut microbiota composition changes with age, behavioral task, and nutrition in the field^24,25^, it would be interesting to validate that variation in the composition of the fecal microbiota mirrors that of the whole gut under such internal and external constraints. Repeated sampling of feces did not affect bees’ survival in a previous study where feces were sampled once per week across three weeks^5^. The possibility of non-invasively monitoring gut microbiota composition via fecal matter collection will help identify the sources of variation in individual gut microbial communities and link this variation to concomitant changes in host phenotypes^20^. This will facilitate longitudinal field studies on natural populations of honey bees and wild bee species to further characterize ecological and evolutionary processes shaping host-microbe interaction^20,26^. Feces sampling might also be used as a tool to assess pathogen loads in the gut. Copley and colleagues found that the gut parasites *Nosema apis* and *Nosema ceranae* could be detected in the feces of contaminated honey bees^27^. However, the correlation between pathogen abundance in the feces and the gut still needs to be uncovered.

The presence of all core bacterial phylotypes in the gut of bees transplanted with a fecal inoculum confirms that the core gut microbiota can be acquired via ingestion of fecal matter, as suggested by previous studies^11,12^. While newly emerged honey bees probably encounter feces on the contaminated hive material^12^, coprophagy, a behavior consisting of feces consumption, has not been described in this insect. It is, however, common in gregarious and social insects and allows transmission of the gut microbiota between overlapping generations^28^. Insects may also benefit from the anti-microbial properties of feces via this behaviour^29^. For instance, fecal transplantation in newly emerged bumblebees led to the development of a gut microbiota similar to that of bumblebees from the donor and protected them against the gut parasite *Crithidia bombi*^29,30^. Our results push for the use of fecal transplantation to study the effect of gut microbiota transmission on microbial communities and host phenotypes with a more ecological approach compared to the currently used inoculation of gut homogenate. Given the volume of feces that can be collected from a single bee without altering its physiology (4.8 ± 2.0 μl on average^5^), one can reasonably expect to inoculate at least four MF bees with feces from a single donor in future experiments. Further dilution of fecal material would likely still allow successful seeding of the gut microbiota of MF bees and would enable inoculation of more individuals.

Finally, we also found that while the bacterial communities in the feces and gut were more similar when originating from the same donor bee, such pairing did not transfer to generation no. 2 when analyzing the gut microbiota of bees inoculated with paired samples. Furthermore, paired samples across generations (*i.e.* gut of a bee from generation no. 2 and its inoculum) did not show greater similarity in microbiota composition than unpaired samples. Such decoupling of community structure across generations, likely suggests that community assembly mechanisms and the rearing environment play a greater role than the inoculum source in determining the final composition of the gut communities. Rearing bees of generation no. 2 in cages where social interaction, in particular trophallaxis events, and coprophagy are possible might have influenced the establishment of the bacterial community in the gut of inoculated bees, additionally to other known mechanisms affecting community assembly (*e.g.* interactions between different bacterial community members and between bacteria and the host)^31^.

In conclusion, our study not only confirms the hypothesis that the honey bee gut microbiota can be transmitted through contact with fecal matter but also opens doors toward longitudinal analyses of individual variation in gut microbiota composition. Feces sampling is a non-invasive method that will reduce the number of animals killed for experimental purposes. This is particularly critical for the study of endangered bee species, or species that are rare or difficult to maintain in laboratory settings. Future studies should yet confirm that feces are a good proxy for gut microbiota composition in other bee species. Fecal transplantation will offer unprecedented opportunities for studying host-microbe interactions in a non-destructive and ecologically relevant manner, as already done in humans and laboratory rodents^19,32^.

## Methods

### Honey bee rearing and gut colonization

Microbiota-free (MF) honey bees *Apis mellifera carnica* were obtained from four hives located at the University of Lausanne (VD, Switzerland), as previously described^33^. Briefly, mature pupae were transferred from capped brood frames to a sterile plastic box for each visited hive and they were kept in a dark incubator for 3 days (35 ℃ with 75 % humidity). Adult bees emerging in such laboratory conditions are MF as their gut is free of any symbionts. Bees had unlimited access to a source of sterile 1:1 (w/v) sucrose solution for the duration of the experiment.

On the third day, five adult nurse honey bees were collected from each of the four original hives (Figure 1). They were stunned using CO_2_ and immobilized on ice at 4 ℃, and their feces and guts were sampled as described previously^5,16^. Two volumes of 2 μl were collected from each fecal sample and diluted 1:10 (v/v) in either sterile PBS or with 1:1 (v/v) PBS:sucrose solution. Gut samples were homogenized in 1 ml of sterile PBS in bead-beating tubes containing zirconia beads using a FastPrep-25 5G apparatus (MP Biomedicals) set at 6 m s^-1^ for 30 sec. Homogenized gut samples were then diluted 1:10 (v/v) to a final volume of 100 μl with 1:1 (v/v) PBS:sucrose solution. The PBS-diluted gut and fecal samples were stored at −80 ℃ for further DNA extraction. They constitute the samples of generation no. 1 (Fig. 1). Feces and gut samples resuspended in PBS-sucrose solution were immediately used for the colonization of MF honey bees.

Gut colonization was carried out by individually pipette-feeding MF bees 5 μl of either diluted feces or gut homogenate, which were sourced from bees originating from the same hive. Additionally, each pair of bees colonized with feces or gut sampled from the same nurse bee were marked by a unique color mark painted on their thorax. It enabled the matching of individuals between generations. Colonized bees were kept in groups of 5 individuals in separate sterile cup cages according to their inoculum and hive of origin at 32 ℃ with 75 % humidity. Bees had access to a sterile sucrose solution and pollen sterilized by gamma-irradiation *ad libitum*.

After 7 days, honey bees were immobilized on ice at 4 ℃, sacrificed and their guts were dissected. Gut samples were homogenized as described above and stored at −80 ℃ for further DNA extraction. They were considered samples of generation no. 2 (Fig. 1).

### DNA extraction

DNA was extracted from the feces of bees from generation no. 1 (Fig. 1) and from the gut of bees from generation no. 1 and 2 (Fig. 1). Homogenized gut tissues were thawed on ice, and 478 μl of those were used for the DNA extraction procedure. The fecal samples were thawed on ice and diluted by mixing 15 μl of feces with additional sterile PBS, to a final volume of 478 μl. For the following steps, both diluted feces and homogenized guts were treated in the same way.

Each sample received 20 μl of 20 mg ml^-1^ proteinase K and 2 μl of s-mercaptoethanol, resulting in 500 μl of source material. Samples were then diluted 2:1 (v/v) with 2X hexadecyltrimethylammonium bromide (CTAB), mixed by bead-beating with glass and zirconia beads using the FastPrep-25 5G set at 6 m s^-1^ for 30 sec and incubated at 56 ℃ for 1 h. Samples were mixed with 750 μl of phenol-chloroform-isoamyl alcohol (PCI; ratio 25:24:1; pH = 8), and centrifuged at room temperature for 10 min at 16,000 *g.* The upper aqueous layer was transferred to a new tube with 500 μl of chloroform and mixed by vortexing. Samples were centrifuged again at room temperature for 10 min at 16,000 *g.* The upper aqueous layer was mixed with 900 μl of cold 100% ethanol and incubated overnight at −20 ℃ to allow for DNA precipitation. Samples were centrifuged at 4 ℃ for 30 min at 16,000 *g* and the supernatant was discarded. DNA pellets were gently washed with 70% ice-cold ethanol, before being centrifuged again at 4 ℃ for 15 min at 16,000 *g.* The supernatant was discarded, and the remaining ethanol was evaporated at room temperature for approximately 10 min. Dried DNA pellets were dissolved in 50 μl of nuclease-free water by incubation at 64℃ for 10 min. Purification of the extracted DNA using CleanNGS magnetic beads (CleanNA) was automated with an Opentrons OT-2 pipetting robot. Briefly, DNA extracts were incubated with 25 µL of NGS beads at room temperature for 10 min. A magnet was involved to attract the beads and attached DNA at the bottom and clear the supernatant. Beads were rinsed twice with 110 µL of ethanol (80%) and left to dry at room temperature for 10min. The obtained purified DNA extracts were resuspended in 45 µL of Tris-HCl buffer (5 μM; pH 8) and stored at −20 ℃. One sample from hive ten was lost during the DNA extraction process.

### 16S rRNA amplicon-sequencing

The extracted DNA was used as a template for 16S rRNA amplicon sequencing following the Illumina metagenomic sequencing official guidelines. Briefly, the 16S rRNA gene V4 region was amplified with the primers 515F (5’-TCGTCGGCAGCGTCAGATGTGTATAAGAGACAGGTGCCAGCMGCCGCGGTAA-3’) and 806R (5’-GTCTCGTGGGCTCGGAGATGTGTATAAGAGACAGGGACTACHVGGGTWTCTAAT-3’), using a high-fidelity polymerase (Phanta Max, Vazyme). PCR products were purified using CleanNGS magnetic beads (CleanNA) in a ratio of 0.8:1 beads to PCR product. Index PCR was performed using Illumina Nextera Index Kit v2 adapters and resulting amplicons were purified again using CleanNGS magnetic beads. PCR products were purified once more using CleanNGS beads in a ratio of 1.12:1 beads to PCR product. Sample concentrations were normalized based on PicoGreen (Invitrogen) quantification and pooled together. Short-read amplicon sequencing was carried out with an Illumina MiSeq sequencer at the Genomic Technology Facility of the University of Lausanne (Switzerland), producing 2 x 250-bp paired-end reads *via* 150 cycles. Negative controls of DNA extraction and PCR amplification were also sequenced for reference.

### Microbial community structure analyses

The bacterial communities present in fecal and gut samples were determined based on analysis of Illumina sequencing, as previously described^8^. Briefly, raw sequencing data were pre-processed by clipping the primer sequences from all reads using Cutadapt^34^ (version 4.2 with Python version 3.11.2).

Sequencing data were then processed following the Divisive Amplicon Denoising Algorithm 2 pipeline^35^ (DADA2; version 3.16) run with R (version 4.2.2). The end of sequences with low quality were further trimmed after 232 and 231 bp, for forward and reverse reads, respectively.

The resulting reads were denoised using the core sample inference algorithm of DADA2, based on error rate learning determined by analyzing 3^8^ minimum numbers of total bases from samples picked at random (‘nbases’ and ‘randomize’ arguments), and paired-end sequences were merged. Unique sequences outside the 250:255-bp range were removed alongside chimeras. The obtained amplicon-sequence variants (ASVs) were classified using the SILVA reference database (version 138.1)^36^. The taxonomic classification was complemented *via* Blast searches to further discriminate ASVs identified as the genus *Lactobacillus* as either the core phylotypes Firm-5 and Firm-4 of the bee gut microbiota or other non-core *Lactobacillus* species. The dataset was cleaned using Phyloseq^37^ (version 1.42.0) by removing any unclassified and eukaryotes ASVs. Lastly, the R package Decontam^38^ (version 1.18.0) was used to remove contaminants based on prevalence and frequency methods.

### Bacterial load quantification by qPCR

Bacterial loads in the gut and feces samples were determined from quantitative PCRs (qPCRs), as previously described^33^. Briefly, universal primers of the 16S rRNA gene were used to determine bacterial load (forward: 5’-AGGATTAGATACCCTGGTAGTCC-3’; reverse: 5’-YCGTACTCCCCAGGCGG-3’) and primers specific to the *Actin* gene of *A. mellifera* were employed as control of sample quality (forward: 5’-TGCCAACACTGTCCTTTCTG-3’; reverse: 5’-AGAATTGACCCACCAATCCA-3’). Corresponding standard curves were generated using serial dilutions of plasmids bearing the target sequences for the 16S rRNA and *Actin* genes.

Purified DNA was used as a template for qPCR reactions by mixing 1 μl of DNA to 5 μl of 2X SYBR Select Master Mix (ThermoFisher), 3.6 μl of nuclease-free water, and 0.2 μl of each appropriate 5 μM primers. Amplification reactions were performed with a QuantStudio 5 real-time PCR machine (ThermoFisher), with the following thermal cycling conditions: 50 ℃ for 2 min and 95℃ for 2 min for denaturation of DNA, followed by 40 amplification cycles consisting of 95 ℃ for 15 sec and 60 ℃ for 1 min. Each reaction was performed in triplicate. The quantification of gene copy numbers was performed following a published detailed protocol^33^. The slope of the standard curves for each target (*i.e.* universal 16S rRNA gene and *actin*) was used to calculate the primer efficiencies (*E*) according to the equation: *E* = 10(–1/slope). The copy number *n* in 1 µL of DNA was obtained using the formula *n* = *E* ^(intercept-Cq)^. This number was multiplied by the elution volume of the DNA extract to obtain the copy number per gut. Finally, the bacterial 16S rRNA gene copy number was normalized for each sample by dividing it by the corresponding *actin* copy number and multiplying by the median of *actin* copy numbers across all samples.

### Statistical analyses

All statistical analyses were performed in R (version 4.2.2). Absolute abundances of each ASV in each sample were calculated by multiplying their proportion by the normalized 16S rRNA gene copy number measured by qPCR. Measures of α diversity (Shannon and Simpson metrics) were obtained with the Phyloseq package^37^ (version 1.42.0). Their normal distribution and homoscedasticity were assessed using a Shapiro-Wilk test and a Bartlett test respectively. For normally distributed and homoscedastic data, differences in α diversity metrics between sample types were tested using paired t-tests, otherwise they were analyzed with two-sided Wilcoxon matched-pairs tests. Difference in community structure was assessed based on Bray-Curtis dissimilarities (Phyloseq) using a Adonis and Permutation test (vegan^39^; version 2.6-4). Estimation of correlation between sample pairs was done using Procuste and Mantel tests based on the Pearson correlation method (ade4^40^; version 1.7.22 and vegan). ASVs with significant differences in their relative abundances between sample types in generation no. 1 were determined using the DESeq2 package^41^ (version 1.38.3).

## Acknowledgements

We thank Florian Zoppi for his continuous support in the laboratory. We also thank the Genomic Technologies Facility of the University of Lausanne, Switzerland, for performing Illumina sequencing. This work was supported by the Marie Skłodowska-Curie fellowships HarmHoney (grant no. 892574, awarded to A.Ca.) and BRAIN (grant no. 797113, awarded to J.L.), the NCCR Microbiomes (National Centre of Competence in Research), funded by the Swiss National Science Foundation (SNSF, grant no. 180575), and an ERC Starting Grant (MicroBeeOme, grant no. 714804) and SNSF Consolidator grant (grant no. TMCG-3_213860) both awarded to P.E..

## Author contributions

The original idea for this manuscript emerged from discussions between A.Ca., A.Ch. and J.L.. A.Ca. and A.Ch. conceived the study, designed experiments, carried out bee experiments, and DNA extractions. N.N. performed the purification of DNA samples. L.K. performed qPCR experiments. A.Ch. and J.L. performed AmpliSeq libraries preparations. J.L. and A.Ch. analyzed the amplicon sequencing data and quantitative PCR data with assistance from A.Ca.. A.Ch. plotted the graphs. Y.S. and P.E. supervised the project. A.Ca. and A.Ch. drafted the manuscript. All authors edited subsequent drafts.

## Supplementary material

**Supplementary Figure 1.**
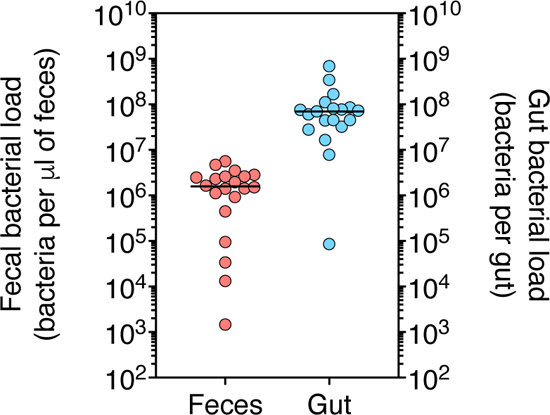
Bacterial loads measured as copies of the 16S rRNA gene in the fecal and gut samples of bees from generation no. 1.

**Supplementary Figure 2.**
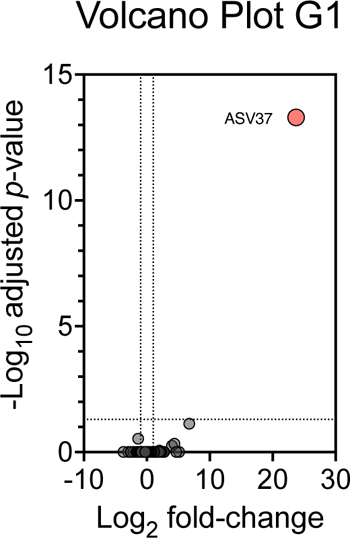
Volcano plot presenting significance vs. fold-change based on relative abundances of all amplicon sequence variants (ASVs) in the gut compared to the fecal samples of nurse bees from generation no. 1. The colored ASV was significantly different in DESeq2 analyses (FDR-corrected *P*<0.05).

**Supplementary Figure 3.**
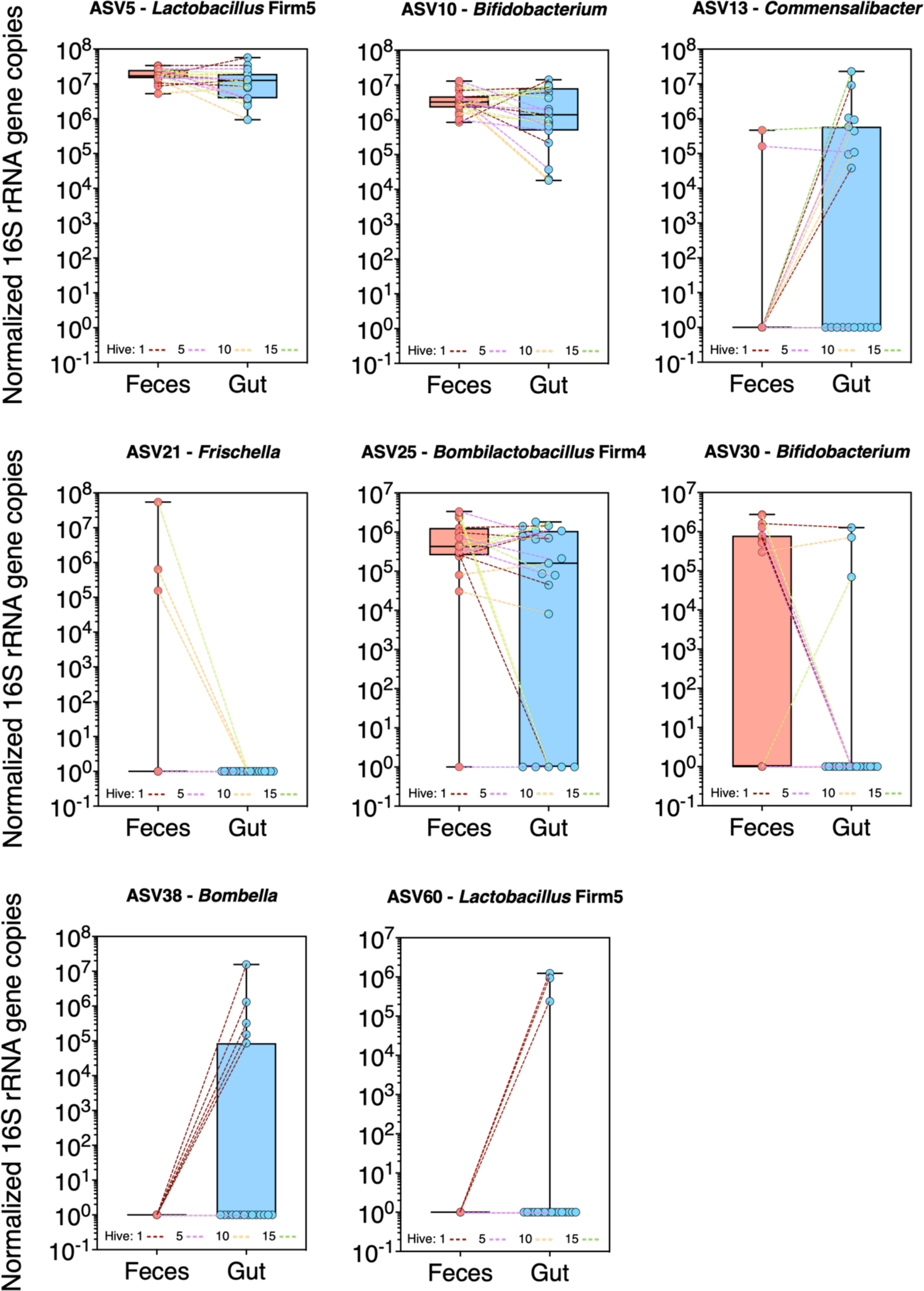
Significantly different 16S rRNA gene copies of various ASVs in the gut of bees inoculated with either feces or gut homogenate.

**Supplementary Figure 4.**
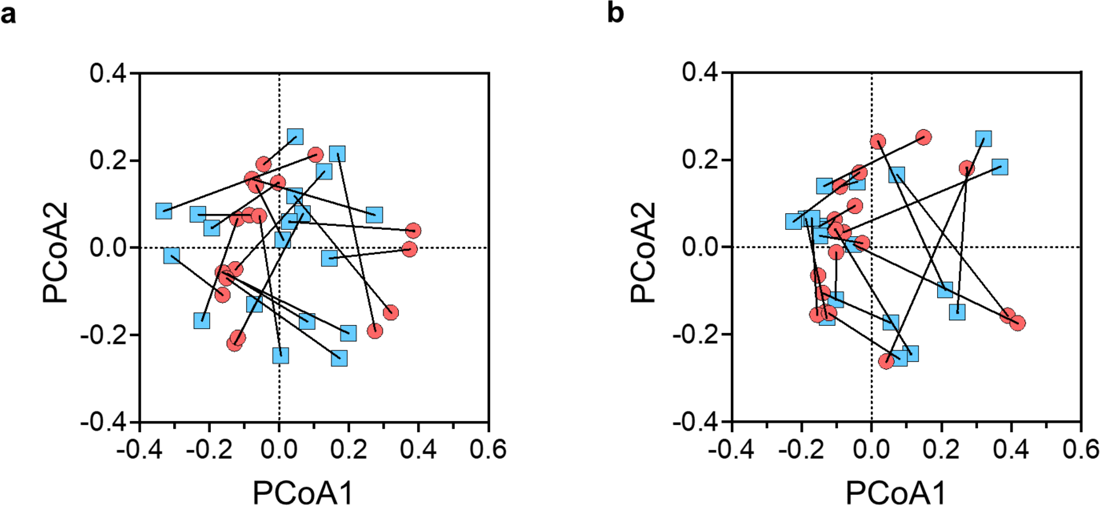
Procrustes analysis of relative ASVs abundances in the fecal (a) or gut (b) samples of bees from generation no. 1 (red) against the gut of bees from generation no. 2 (blue) was obtained from PCoA and revealed no significant agreement of comparison.

